# Myocardial B cells have specific gene expression and predicted interactions in Dilated Cardiomyopathy and Arrhythmogenic Right Ventricular Cardiomyopathy

**DOI:** 10.1101/2023.09.21.558902

**Authors:** Kevin C. Bermea, Carolina Duque, Charles D. Cohen, Aashik Bhalodia, Sylvie Rousseau, Jana Lovell, Marcelle Dina Zita, Monica R. Mugnier, Luigi Adamo

**Affiliations:** Division of Cardiology, Department of Medicine, Johns Hopkins University School of Medicine, Baltimore, Maryland; Department of Pathology, Johns Hopkins University School of Medicine, Baltimore, Maryland; Department of Molecular Microbiology and Immunology, Johns Hopkins Bloomberg School of Public Health, Baltimore, Maryland, USA

## Abstract

**Introduction:** Growing evidence from animal models indicates that the myocardium hosts a population of B cells that play a role in the development of cardiomyopathy. However, there is minimal data on human myocardial B cells in the context of cardiomyopathy.

**Methods:** We integrated single-cell and single-nuclei datasets from 45 healthy human hearts, 70 hearts with dilated cardiomyopathy (DCM), and 8 hearts with Arrhythmogenic Right Ventricular Cardiomyopathy (ARVC). Interactions between B cells and other cell types were investigated using the CellChat Package. Differential gene expression analysis comparing B cells across conditions was performed using DESeq2. Pathway analysis was performed using Ingenuity, KEGG, and GO pathways analysis.

**Results:** We identified 1,100 B cells, including naive B cells and plasma cells. B cells showed an extensive network of interactions within the healthy myocardium that included outgoing signaling to macrophages, T cells, endothelial cells, and pericytes, and incoming signaling from endothelial cells, pericytes, and fibroblasts. This niche relied on ECM-receptor, contact, and paracrine interaction; and changed significantly in the context of cardiomyopathy, displaying disease-specific features. Differential gene expression analysis showed that in the context of DCM both naive and plasma B cells upregulated several pathways related to immune activation, including upregulation of oxidative phosphorylation, upregulation of leukocyte extravasation, and, in naive B cells, antigen presentation.

**Discussion:** The human myocardium contains naive B cells and plasma cells, integrated into a diverse and dynamic niche that has distinctive features in healthy myocardium, DCM, and ARVC. Naive myocardial-associated B cells likely contribute to the pathogenesis of human DCM.

## INTRODUCTION

The immune system has been highlighted as a key potential player in the development of heart failure (1–4). The role of macrophages, monocytes, and T cells in the context of heart failure has been extensively studied in both human and animal models of cardiomyopathy(1). In recent years, there has been increasing interest in understanding the role that B cells play in cardiomyopathy, but the available data is limited(5).

Studies on murine naive hearts have shown that B cells continuously patrol the heart and circulate between the heart and spleen along the cardio-splenic axis (4, 6–9). B cells residing in the murine myocardium are therefore mainly part of a pool of circulating B cells that transiently adheres to the microvascular endothelium (6, 10) . Intriguingly, a minor subset of myocardial B cells, in both mice and humans, enters the interstitial spaces, indicating a nuanced distribution of these cell types within the cardiac tissue (6, 8). Data from murine models suggests that B cells interact with macrophages in the naive myocardium(11) and play a role in the context of myocardial adaptation to injury and heart failure(6, 10, 12–16). Yet, data about the role that B cells play in the diseased human heart is lacking.

To address this gap in knowledge we performed a focused analysis of single cells and single nuclei datasets from healthy controls and patients with two different forms of cardiomyopathy: dilated cardiomyopathy (DCM) and Arrythmogenic Cardiomyopathy. DCM is a form of cardiomyopathy characterized by reduced left ventricular ejection fraction and increased left ventricular diameters. DCM can have various etiologies that lead to a common phenotype (2). ARVC is a genetic disorder that is typically due to mutations in desmosomal proteins that lead to the replacement of the right ventricular myocardium with ‘fibrofatty’ (mix of fibrous and fatty components) tissue, resulting in dyskinesia of the right ventricle, arrhythmias and often also reduced left ventricular ejection fraction (17–21). We first investigated interactions between B cells and other cell types to gain insight into the biological niche of myocardial B cells, in health and disease. We then performed differential gene expression analysis to gain additional insight into the potential role that B cells might play in various forms of cardiomyopathy. (Figure 1A)

**Figure 1.**
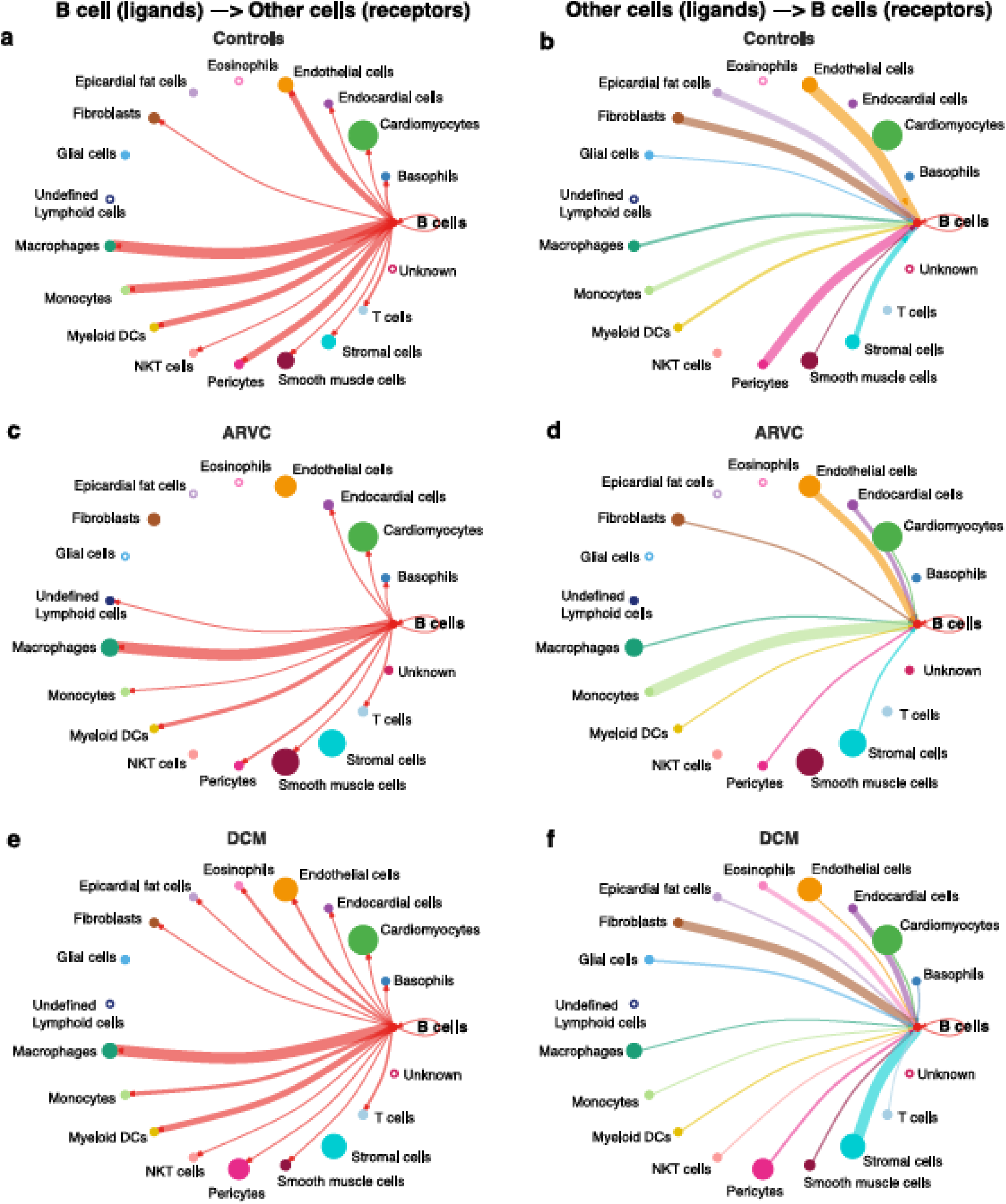
Graphic depicting the *in silico* workflow UMAP and plots generated. **a**): two human heart single cell datasets were downloaded, loaded into R and analyzed using Seurat. Standard quality control and clustering methods were first performed. B cells were then identified using ScType, and ScTransform was used to normalize, scale, and integrate the datasets. Finally, differentially expressed genes were identified and pathway analysis as well as cell-cell communication analysis was performed. Panel b shows a UMAP with the subset of cells identified as B cells that were subsetted from the integrated data and sub-classified into B cell subtypes. Cells with an identity that went unresolved were classified as “unknown” and excluded from the analysis. Panels **c**-**e** show the UMAPs of the integrated data sets from sc/sn-RNA sequencing cells split by condition, including, control (panel **c**), ARVC (panel **d**), and DCM (panel **e**).

## METHODS

### Quality-control

Pre-processed single-cell and single-nuclei datasets were obtained from GSE183852(22) of Gene Expression Omnibus (GEO) and Cellxgene website (https://cellxgene.cziscience.com/collections/e75342a8-0f3b-4ec5-8ee1-245a23e0f7cb/private)(23). These datasets were previously published (22) (23) and it was reported that studies were approved by the appropriate institutional and/or national research ethics committee and were performed in accordance with the ethical standards. These datasets include data from 45 healthy control human hearts, 70 hearts with DCM, and 8 with ARVC. Data from one patient with non-compaction cardiomyopathy was included in the datasets but was excluded from the analysis. Data were loaded onto R (v. 4.1.3)(24) and were analyzed using Seurat (v. 4.1.0)(25–28) with default settings unless otherwise stated. Cells were verified to include only cells with unique feature counts between 200 – 15,000 and mitochondrial read counts <5% . Ensemble gene IDs were converted to gene symbols where applicable using biomaRt(29), and only gene IDs that matched between the GEO and Cellxgene datasets were kept for downstream applications.

### B cell identification

Cells classified as lymphocytes by the authors of each dataset were subsetted and counts were normalized using the logNormalize function that takes feature counts for each cell and divides it by the total counts for that cell, then multiplies by a scale factor of 10,000, and natural-log transforms the values. The data were then scaled using the ScaleData function. The 2000 most highly variable features were identified using the FindVariableFeatures function with the variance stabilizing transformation method. Then, the function RunPCA was used, with 50 principal components (PCs) generated. The first 10 PCs were used to find the shared nearest neighbors (SNN) using the FindNeighbours function. Cluster identification was performed using the FindClusters function with a resolution of 0.8 using the Louvain algorithm, and a UMAP was created with the first 10 PCs. Subsequently, the ScType algorithm was employed for the identification of B cells based on known human B cell markers (Table S1)(30, 31).

### B cell Integration and Clustering

For visualization, SCTransform was utilized to normalize, scale, identify variable features, and regress out percent mitochondrial gene expression. SCTransform (v2) was performed using the gamma-Poisson generalized linear model. After SCTransform, 6000 consistently variable features across datasets were selected as integration features using the SelectIntegrationFeatures function. Next, the integration features were subjected to principal component analysis using RunPCA. Integration anchors were identified using the FindIntegrationAnchors function using reciprocal PCA dimensional reduction based on the first 50 PCs, using 10 neighbors (k) when choosing anchors. These anchors were used to integrate the data using the first 50 dimensions for the anchor weighing procedure in the IntegrateData function.

After integration, PCA was performed again, and a UMAP and SNN graphs were generated based on the first 20 principal components. The cluster identification was made using the Louvain algorithm with a clustering resolution of 0.4.

### B cell differential Gene Expression and Pathway analysis

Differential gene expression between each disease condition (DCM and ARVC) and controls was conducted for each B cell subtype cluster utilizing the DESeq2 R package^32^. This pseudobulk analysis incorporated adjustments for experimental modalities, including whether the data originated from single-cell or single-nuclei protocols, as well as the dataset’s source. Read counts were normalized via DESeq2’s median of ratios method. The Wald test was employed for determining differential expression, with p-values adjusted using the Benjamini-Hochberg procedure to control the false discovery rate. Genes with a raw p-value <0.05 and an absolute fold change >1.5 were used for pathway analyses. For Ingenuity Pathway Analysis (IPA)(32), the Ingenuity Knowledge Base reference set was selected with 35 molecules per network, and 25 networks per analysis for the interaction networks. Pathways with p-value < 0.05 and absolute z-score > 1.5 were treated as statistically significant. Gene ontology (GO) enrichment analysis was performed using the enrichGO function of clusterProfiler (v. 4.4.4). Both analyses were performed with a background gene set of all the genes submitted to differential gene enrichment analysis. The enrichGO function applies an over-representation analysis based on a one-sided Fisher’s exact test to the DEGs detected by DESeq2 analyses. Benjamini-Hochberg-adjusted p-values less than 0.05 are considered significantly enriched pathways. Additionally, the same genes were utilized as input for the Kyoto Encyclopedia of Genes and Genomes (KEGG) pathway analysis using the Database for Annotation, Visualization, and Integrated Discovery (DAVID) tool.

### Whole heart integration and cell annotation

Both datasets were split by disease and by sample type (single-cell vs. single-nuclei). Using SCTransform(33, 34), mitochondrial genes were regressed out, data were normalized, transformed, and scaled, and 3000 variable features were identified. Integration of each dataset’s single-nuclei and single-cell data was performed per disease state in the same way as for the B cell clustering described above, except using 3000 integration features, and employing a referenced based integration. The Cellxgene’s single nuclei dataset was used as reference. Non compaction cardiomyopathy was not integrated, since it was only present in one single nuclei dataset. After integration, PCs were regenerated, and an elbow plot was used to determine the number of significant PCs to use for downstream analyses (20 PCs were used for ARVC, 30 PCs were used for DCM and controls). Cell clustering was performed using Seurat’s FindNeighbours and FindClusters using the Louvain algorithm with a resolution of 0.5 for ARVC, and 0.7 for DCM and controls. Clusters were visualized using UMAP dimensional reduction and annotated using ScType(30) with modified cell markers. The full set of markers used to identify various cell types is reported in Table S1. Clusters annotated as immune cell types were subset out, and then SCTransform, integration, clustering, and annotation were performed again on this immune subset as described above, except using a greater clustering resolution of 0.7 for ARVC and 0.9 for DCM and controls. The new immune cell type annotations were added back to the parent dataset. For simplicity, annotations were condensed as follows: Memory/naive/Effector CD8+/CD4+ T cells to “T cells”; Memory/naive B cell or plasma cells to “B cells”; Non-classical, classical and intermediate monocytes to “Monocytes”; Vascular / Lymphatic endothelial cells to “Endothelial cells”; CD8+ NKT-like T cells to “NKT cells”; and Schwann cells to “Glial cells”.

### Cell-cell-communication

Cell-cell communication was determined using CellChat R package (version 1.6.1) and the human ligand-receptor database, CellChatDB(35). The SCTransformed data was used for all disease groups. For each disease group, we subsetted the data to include only known cell-signaling genes included in the CellChat reference database. Next, we determined the over-expressed ligands and receptors per cell type, over-expressed ligand/receptor pairs across cell types, and the communication probability based on the trimean method, following the standard workflow(35). We calculated the aggregated cell-cell communication network for B cells using the aggregateNet function(35). For visualization, the three disease state objects were “lifted” such that each object included the same set of annotated cell populations so that they could subsequently be merged into a single cellchat object for downstream analyses.

## RESULTS

### In silico analysis of integrated myocardial lymphocytes identifies plasma and naive B cells within the healthy and diseased human heart

B cells and plasma cells from Cellxgene and the GEO datasets were identified, subsetted, integrated, and further subclassified using ScType with selected markers for B cell/plasma cell populations. From 45 control samples and 79 cardiomyopathy samples, encompassing 1,100,752 single nuclei and 49.723 single cells (Figure 1A) a total of 1,100 B cells/plasma cells were identified. These cells were subsequently subsetted and integrated using SCTransform. The integrated data was processed as aforementioned in the methods section, yielding 2 major B cell clusters (Figure 1B, Figure S1). We identified one of the major clusters as naive B cells and the other cluster as plasma cells. We also identified some B cells that we could not confidently assign to any specific B cell subtype and we therefore labeled these as “unknown” (Figure. 1B, Figure S1).

### Myocardial B Cells have an extensive network of interactions with other myocardial cell types, that change by disease state

We first investigated communication between B cells and other cell types to define the biological niche of myocardial B cells in health and disease. This this end, the CellChat algorithm was employed to calculate the likelihood of cell-cell communications. First, in order to identify the various cell types in the heart, clusters in the integrated datasets were annotated using scType (Figure 1 C-E). Next, an evaluation of the relative interaction strength of B cells with other heart cells within a cell-cell communication network was performed, based on CellChat’s calculated communication probability (Figure 2; Figure S2). In control human hearts, B cells were predicted to communicate with macrophages, monocytes, myeloid dendritic cells (DCs), pericytes, endothelial cells, and fibroblasts (Figure 2 A-B). In ARVC, the analysis displayed similar interaction patterns as in controls. However, there were more signals from monocytes, endocardial cells, and cardiomyocytes to B cells; there were reduced signals to or from fibroblasts, pericytes, and stromal cells (Figure 2 C-D; Figure S2 A-B). In DCM, we observed increased signaling from B cells to eosinophils, fibroblasts, and epicardial fat cells. Furthermore, we noted augmented signaling from almost all cells, but especially from stromal cells and fibroblast to B cells. Also, reduced signaling from B cells to monocytes, pericytes, and endothelial cells was observed (Figure 2 E-F; Figure S2 C-D).

**Figure 2.**
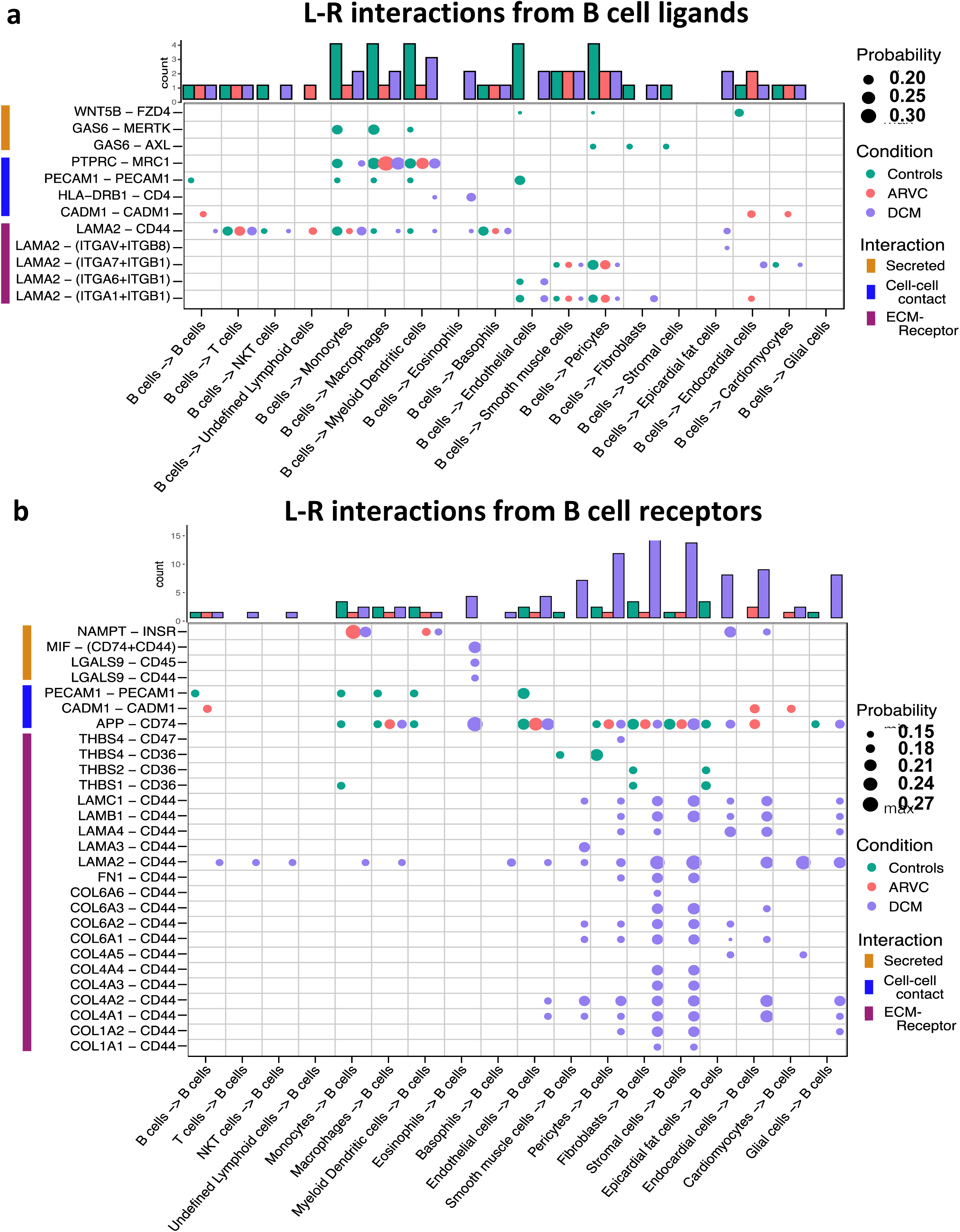
B cell interaction network is notably altered by disease state. Intercellular communications plot of the interactions from B cell ligands to other receptors in **a**) controls, **c**) ARVC, **e**) DCM, and from other cell ligands to B cell receptors in **b**) controls, **d**) ARVC and **f**) DCM. Thickness of the line is relative to the maximum communication probability to or from B cells. Line color represents the source (ligand provider) of the interaction. Size of circle is proportional to the number of cells of that type. Open circles represent cell types that are not detected in the disease state. DC = dendritic cell. Statistical significance determined by CellChat’s permutation test.

To further probe these interactions, the specific B cell ligand-receptor interactions by disease state were visualized (Figure 3). We saw that while some ligand-receptor interactions were conserved across disease states, multiple interactions were significant only in certain disease states. Healthy control hearts were uniquely characterized by B cell signaling to endocardial cells, endothelial, fibroblasts, pericytes, and smooth muscle cells via the Wnt family member 5B (WNT5B) – Frizzled-4 (FZD4), and Growth Arrest Specific 6 (GAS6) – AXL tyrosine kinase (AXL) / MER proto-oncogene, tyrosine kinase (MERTK) pathways (Figure 5A). In healthy control hearts, we also observed that B cell signaling was characterized by PECAM1 homophilic interaction with other B cells, endothelial cells, and myeloid cells, which is important for the diapedesis of immune cells and survival signaling(36) (Figure 3A). Myeloid, endothelial, pericyte, fibroblast, stromal, epicardial fat, and glial cells also signaled to B cells via Amyloid precursor protein (APP) – CD74 and thrombospondin-1/2/4 (THBS1/2/4) – CD36. (Figure 3B).

**Figure 3.**
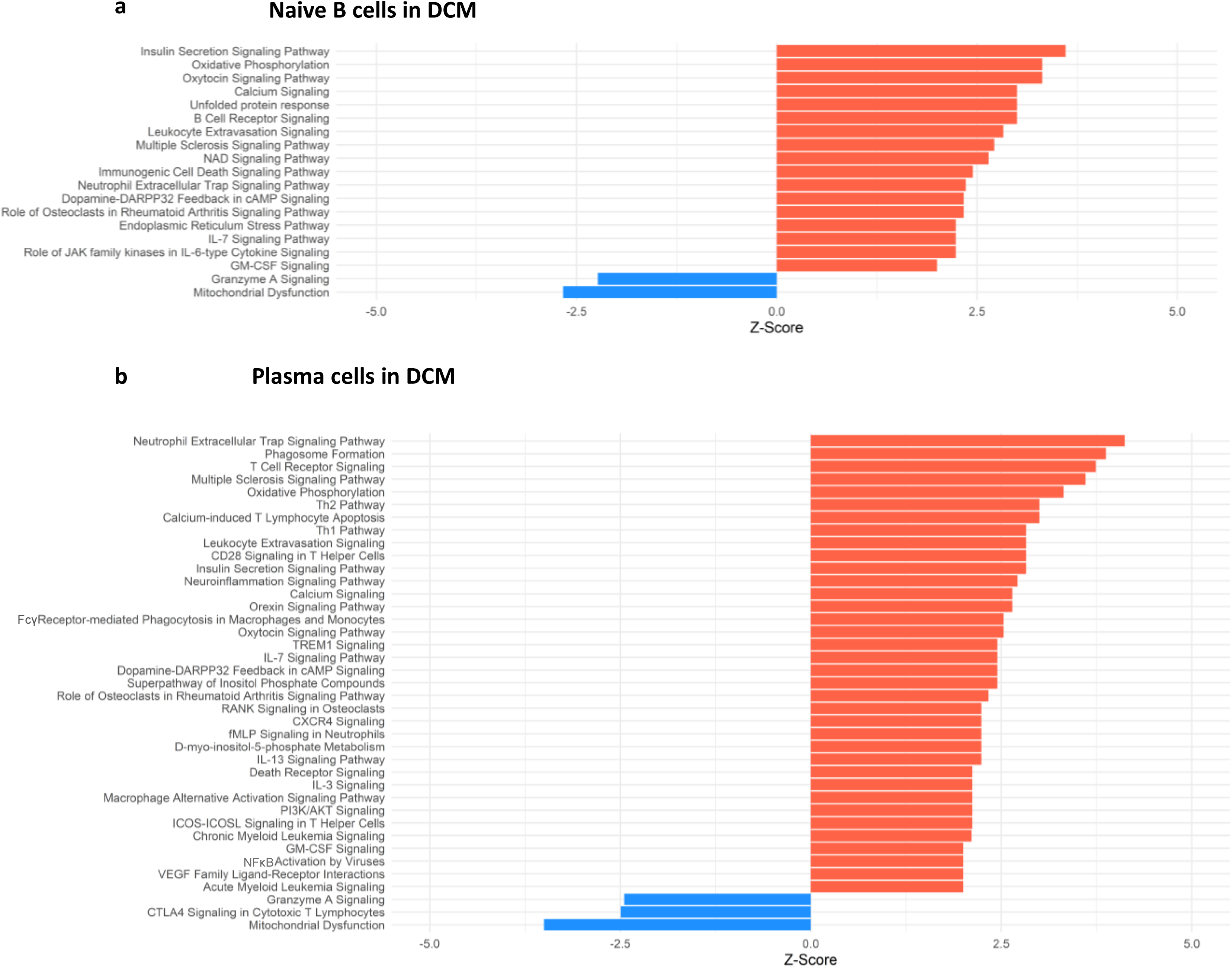
B cell ligand-receptor interactions are significantly altered by disease states. Dot plot of all probable interactions that reached statistical significance. Color represents condition (Controls = turquoise, DCM = purple, ARVC = red). Bar plot represents counts of ligand-receptor interactions for each cell type per disease state. Statistical significance was determined by CellChat’s permutation test. L-R = ligand-receptor, Probability = communication probability as determined by CellChat’s algorithm.

In ARVC, there were overall fewer interactions between B cells and other cell types compared to healthy controls. One cell-cell interaction that was unique to ARVC was the cell-adhesion molecule (CADM1) homophilic interactions between B cells and cardiac cells (Figure 3A). In contrast to ARVC, DCM patients showed considerably increased B cell-cell interactions. B cells signaled to eosinophils and dendritic cells via the MHC-II molecule DR beta 1 (HLA-DRB1) and the CD4 receptor (Figure 3A). In addition, we found that CD44 was a major cell-cell interaction receptor on B cells with extensive interactions with the extracellular matrix (ECM) proteins laminin, collagen, and fibronectin, of various cell types (Figure 3B). Signaling through CD44 has been associated with immune cell migration and activation(37). Eosinophils were also shown to signal to B cells through Macrophage migration inhibitory factor (MIF) – CD44 + CD74 complex which has been associated with B cell chemotaxis and survival(38–40) (Figure 3B). Interestingly, signaling through galectin 9 (LGALS9) – CD45 or CD44 was also observed, which is known to have an inhibitory role on B cell receptor signaling and activation(41, 42) (Figure 3B). Other potential interactions mediated through receptors on B cells included pericyte mediated thrombospondin-4 (THBS4) signaling through CD47 on B cells. B cells were also the target of signalling through CD44 - laminin 2 (LAM2) interactions originating from multiple cell types including epicardial fat cells, fibroblasts, and endocardial cells (Figure 3B).

### Differential gene expression analysis reveals pronounced dysregulation of inflammatory pathways in myocardial naive B Cells and plasma B cells in DCM, but not in ARVC

To further elucidate the biological significance of B cells in human cardiomyopathy, we sought to identify differentially enriched pathways in DCM and ARVC B cell populations. Pseudo-bulk differential gene expression analysis was performed to compare gene expression between disease conditions and controls for each B cell cluster. Few genes were differentially expressed with an adjusted p-value <0.05, so an unadjusted p-value <0.05 and fold change >1.5 were used as cutoffs for including genes in downstream pathway analyses. In DCM, 227 genes met this cutoff in the plasma cells cluster, and 116 in the naive B cells cluster (Table S2 and Table S3). In ARVC, 215 genes and 136 genes met this cutoff in the plasma and naive B cell clusters, respectively (Table S4 and Table S5). These gene lists were submitted to GO, KEGG, and IPA pathway analyses.

In ARVC, the genes differentially expressed in naive B cells and plasma cells showed no enrichment in inflammation-related pathways when analyzed via KEGG pathway (Tables 1 and 2), GO pathways (Table S6), or IPA pathways analysis (Table S7 and Table S8).

**Table 1.**
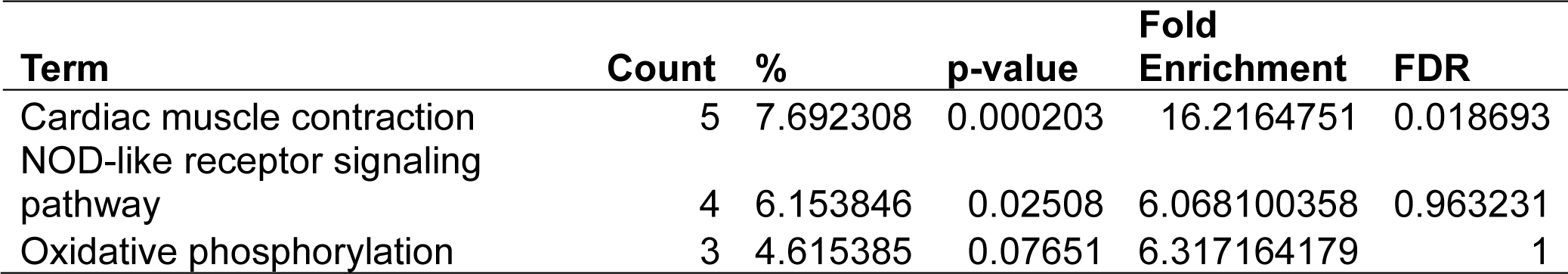
Dysregulated KEGG Pathways Assessed Using DAVID in ARVC’s naive B cells cluster, contrasting ARVC vs control samples data.

| Term | Count | % | p-value | Fold Enrichment | FDR |
| --- | --- | --- | --- | --- | --- |
| Cardiac muscle contraction | 5 | 7.692308 | 0.000203 | 16.2164751 | 0.018693 |
| NOD-like receptor signaling pathway | 4 | 6.153846 | 0.02508 | 6.068100358 | 0.963231 |
| Oxidative phosphorylation | 3 | 4.615385 | 0.07651 | 6.317164179 | 1 |

**Table 2.**
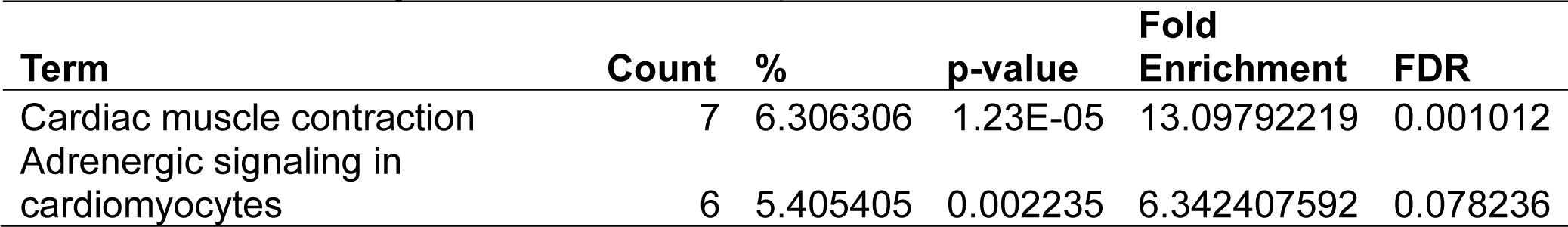
Dysregulated KEGG Pathways Assessed Using DAVID in ARVC’s plasma cells cluster, contrasting ARVC vs control samples data.

| Term | Count | % | p-value | Fold Enrichment | FDR |
| --- | --- | --- | --- | --- | --- |
| Cardiac muscle contraction | 7 | 6.306306 | 1.23E-05 | 13.09792219 | 0.001012 |
| Adrenergic signaling in cardiomyocytes | 6 | 5.405405 | 0.002235 | 6.342407592 | 0.078236 |

Conversely, in DCM both naive B cells and plasma cells showed significant dysregulation of multiple pathways related to the immune response. In the naive B cell cluster, IPA showed upregulation of B Cell Receptor Signaling, Leukocyte Extravasation, and Immunogenic Cell Death Signaling (Figure 4A; Table S9). Additionally, GO enrichment analysis indicated dysregulation of B cell activation, B cell proliferation, antigen process and presentation, and antigen receptor-mediated signaling (Figure S3, Table S10). The top dysregulated pathways related to immune response by KEGG pathway analysis were antigen processing and presentation and B cell receptor signaling pathway (Table 3). When focusing on the plasma cell cluster, in DCM IPA showed upregulation of NFκB Activation by viruses, macrophage alternative activation signaling pathway, IL-3 Signaling, CXCR4 signaling, leukocyte extravasation signaling, T cell receptor signaling, neutrophil extracellular trap signaling pathway, IL-7 signaling pathway, Fcγ receptor-mediated phagocytosis in macrophages and monocytes (Figure 4B; Table S11). GO enrichment analysis showed dysregulation of several pathways that involve transcript processing and protein synthesis (Fig. S3; Table S12). KEGG pathway analysis showed dysregulation in antigen processing and presentation and B cell receptor signaling pathways (Table 4).

**Figure 4.**
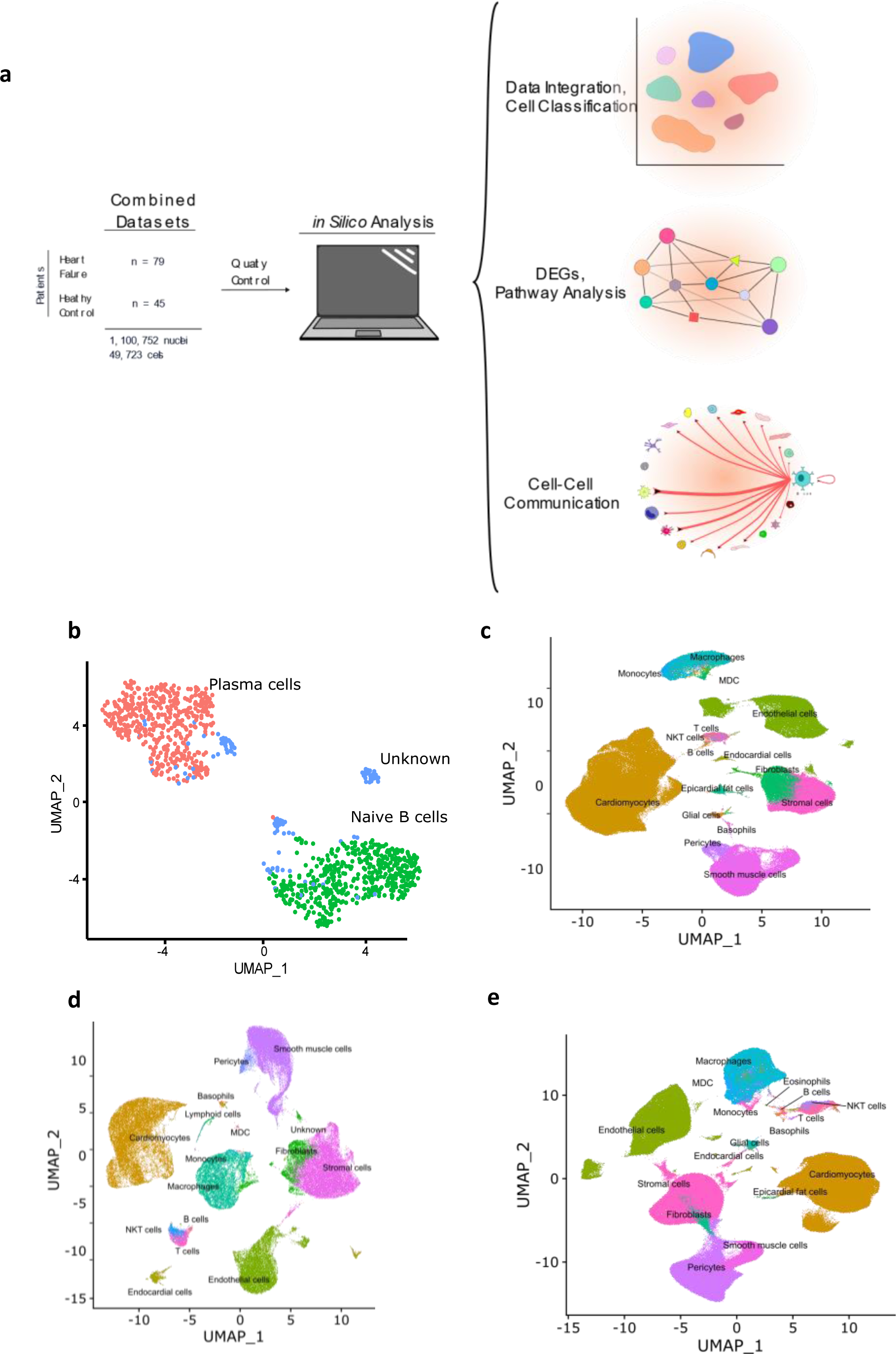
Dysregulated Pathways Identified using IPA in both Naive B cells and Plasma cells clusters from DCM samples as compared to controls. The bar plots depict the dysregulated pathways identified by Ingenuity Pathway Analysis (IPA). **a**) Dysregulated Ingenuity Canonical Pathways in the Plasma cells cluster. **b**) Dysregulated Ingenuity Canonical Pathways in the Naive B cells cluster. The plots show pathways involving immune system response and metabolism. The pathways shown here meet the statistical criteria of a p-value below 0.05 and an absolute z-score exceeding 1.5. ARVC data did not show any relevant pathway on IPA based on the defined threshold criteria.

**Table 3.**
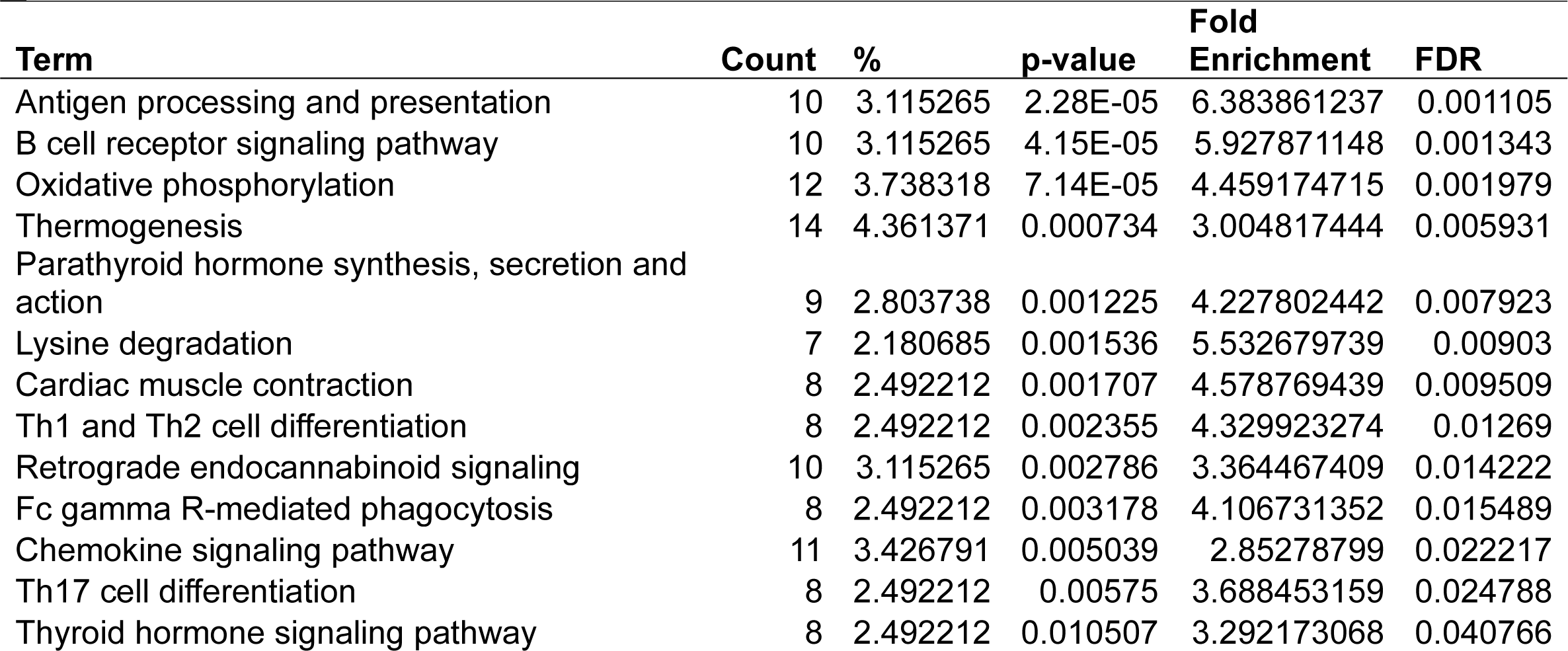
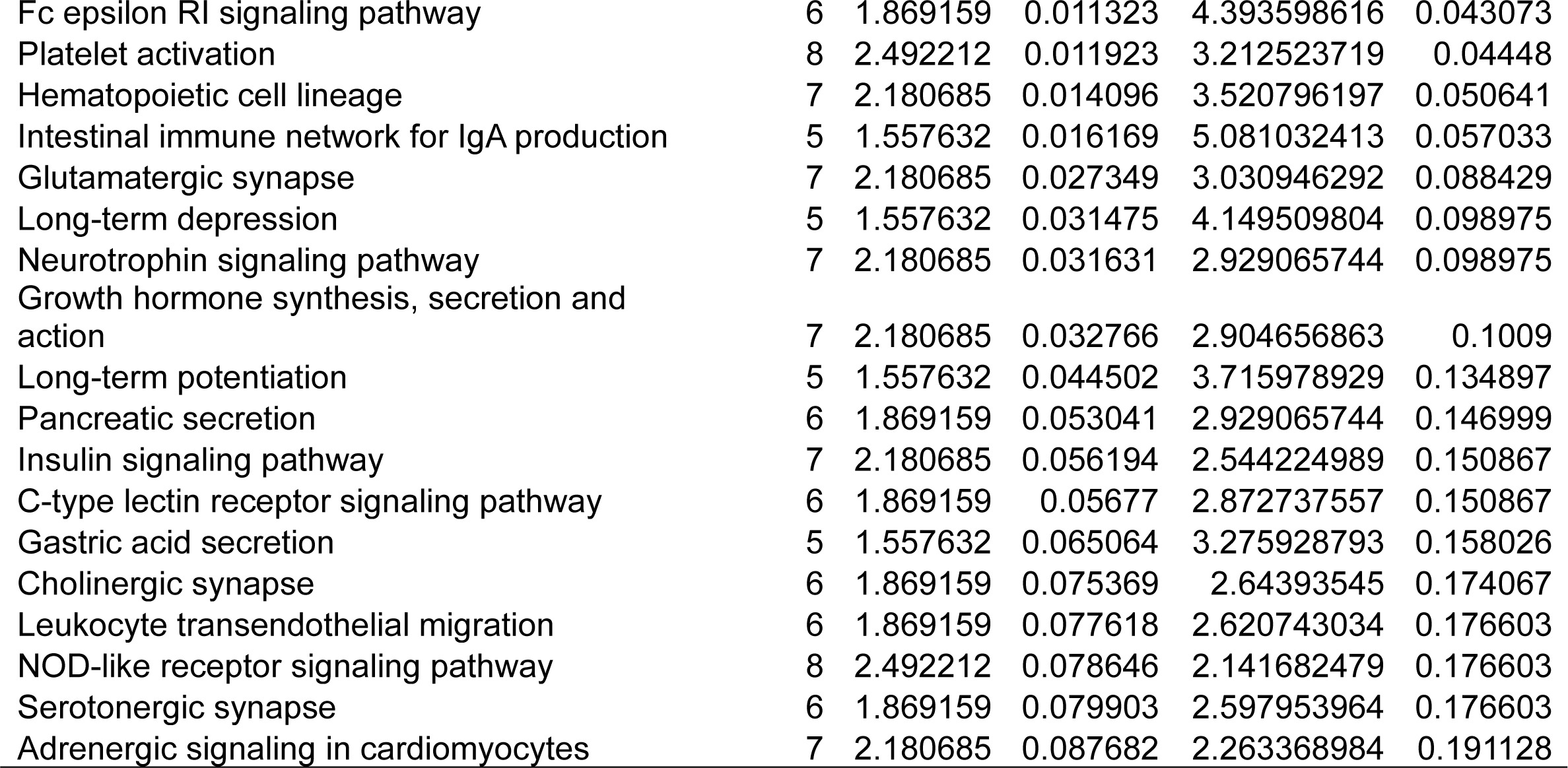
Dysregulated KEGG Pathways Assessed Using DAVID in DCM’s naive B cells cluster, contrasting DCM vs control samples data.

**Table 4.** Dysregulated KEGG Pathways Assessed Using DAVID in DCM’s plasma cells cluster, contrasting DCM vs control samples data.

| Term | Count | % | p-value | Fold Enrichment | FDR |
| --- | --- | --- | --- | --- | --- |
| Cardiac muscle contraction | 12 | 2.348337 | 2.93E-05 | 4.885297937 | 0.003383 |
| Lysine degradation | 9 | 1.761252 | 0.000354 | 5.059772863 | 0.016338 |
| Thermogenesis | 16 | 3.131115 | 0.00225 | 2.442648968 | 0.047242 |
| Oxidative phosphorylation | 11 | 2.152642 | 0.004421 | 2.907481421 | 0.068073 |
| Thyroid hormone signaling pathway | 10 | 1.956947 | 0.006941 | 2.927141326 | 0.094316 |
| Antigen processing and presentation | 7 | 1.369863 | 0.022132 | 3.17857526 | 0.213023 |
| Adrenergic signaling in cardiomyocytes | 10 | 1.956947 | 0.029497 | 2.299896756 | 0.243349 |
| Growth hormone synthesis, secretion and action | 8 | 1.565558 | 0.052089 | 2.361227336 | 0.325206 |
| Parathyroid hormone synthesis, secretion and action | 7 | 1.369863 | 0.077804 | 2.338951607 | 0.438358 |
| Biosynthesis of nucleotide sugars | 4 | 0.782779 | 0.0848 | 3.829017302 | 0.4607 |
| B cell receptor signaling pathway | 6 | 1.174168 | 0.087256 | 2.529886432 | 0.4607 |
| Estrogen signaling pathway | 8 | 1.565558 | 0.093866 | 2.053241162 | 0.481846 |

## DISCUSSION

We present an analysis of human myocardial single cell and single nuclei datasets focused on the analysis of the biological niche of myocardial B cells, in the healthy and diseased heart. We found that the human myocardium harbors naive B cells and plasma cells (Figure 1, Figure S1) that interact with multiple cell types, especially macrophages, monocytes, endothelial cells, pericytes, and fibroblasts (Figure 2). We found that the rich network of interactions of myocardial B cells is altered in the context of cardiomyopathy, with disease-specific features (Figure 2-3, Figure S2). This is reflected by disease-specific changes in gene expression of both naive B cells and plasma cells (Figure 4, Figure S3). These findings expand our understanding of the biology of myocardial B cells and suggest that B cells might play a role in the pathogenesis of specific types of human cardiomyopathies.

Initial studies in murine models identified myocardial B cells as naive B cells and B1 cells(6, 8, 16). Follow-up studies raised the possibility that the myocardium could host several other B cell types including multiple types of follicular B cells, germinal center cells, and marginal zone B cells(15). Data on human myocardial B cells is limited. A study based on analysis of histological sections from multiple non-failing hearts and single-cell sequencing data from 14 healthy human hearts identified a small population of naive B cells with a gene expression signature similar to that observed in rodent studies, as well as plasma cells(43). Plasma cells were also recently described in a study that performed tissue transcriptomic-based analysis of the human myocardium(44). Our findings corroborate the notion that the human heart harbors naive B cells and plasma cells.

There is minimal to no data on the biological niche of human myocardial B cells. Our focused analyses indicate that, in the naive heart, myocardial-associated B cells interact with multiple cell types, both sending signals to other cells and receiving signals from other cells. In the naive heart, B cells appear to have the strongest outgoing communication with macrophages (Fig.2A). This is in line with murine data suggesting that B cells modulate the expression of specific surface markers on resident myocardial macrophages(11). However, B cells appeared to have an extensive network of communication with multiple different cell types (Figure 2A-2B). This is remarkable but, to some extent, not completely unexpected considering that studies in rodents have shown that young mice with congenital B cell deficiency present changes in myocardial structure and function when compared to syngenic, age/sex-matched controls(6).

We found that the network of interactions between B cells and other myocardial cell types changes in the context of cardiomyopathy, with disease-specific features (Fig 2 A-F and Fig 3). This is arguably the most important finding of our study as it suggests that myocardial B cells play a specific role in specific pathological conditions. Notably, the Wnt (WNT5B) – Frizzled (FZD4), GAS6–AXL/MERTK pathways, and *PECAM1* homophilic interactions were prevalent in the healthy state but were lost in disease states (Figure 3A). These pathways are associated with proliferation, growth, and survival signaling(36, 45, 46). This may therefore suggest that B cells play a role in maintaining the cardiac architecture in normal states, a hypothesis that is in line with previously referenced observations in murine models that connect congential B cell deficiency with alterations in myocardial structure(6). PECAM1 homophilic interaction of B cells with endothelial cells is also consistent with the role of PECAM1 in various stages of the extravasation of immune cells, potentially mediating B cell entry into the myocardial interstitiun(47). *PECAM1* has been demonstrated to be crucial for the process of diapedesis and movement to the sites of inflammation(48). In the healthy heart, other cells signaled to B cells largely through the thrombospondin 1, 2 and 4 – CD36 pathway (Figure 3B), which has been shown to be essential in maintaining B cell metabolic function and activation potential(49). These signals are lost in the context of ARVC and DCM. Taken together, these observations suggest that myocardial B cells might play a role in myocardial homeostasis and at the same time receive within the myocardium specific signals that promote their survival and readiness to respond to pathogenic stimuli. These signals might be turned off once a specific B cell response has been triggered and B cells switch from a “homeostatic function” to a “response function”.

The biological niche of B cells, defined by their network of cell-cell interactions, showed disease-specific features. ARVC was overall characterized by a significant reduction in B cell interactions with most other cells (Figure 2 C-D and Figure 3). However, we noted increased interaction strength with macrophages, monocytes, cardiomyocytes, and endocardial cells compared to healthy controls (Figure 2D). These increased interactions were in part due to *CADM1* homophilic interactions (Figure 3). *CADM1* is a cell-cell adhesion molecule that activates the PI3K pathway(50), which is also activated by the B cell receptor(51). The significance of this finding and its relevance in ARVC remains unclear. Conversely to ARVC, our results indicate that B cell interactions are greatly increased in DCM relative to controls. This is consistent with findings that B cell numbers are also increased in the peripheral blood of DCM patients(52) and with other clinical observations that suggest a pathogenic role of B cells in DCM(53). Most notably, we found that in DCM B cells had considerably greater and stronger interactions with fibroblasts, epicardial fat cells, eosinophils, and stromal cells compared to controls (Figure 2F). An increase in interactions with epicardial fat cells is consistent with prior findings suggesting a potential role of epicardial fat-associated B cells in the pathogenesis of ischemic dilated cardiomyopathy(54). We found that in DCM the majority of the communication between B cells and fibroblasts, epicardial fat cells, and other stromal cells was mediated by the interaction of extra-cellular matrix (ECM) proteins such as laminin, collagen, and fibronectin with CD44 (Figure 3B). CD44 is known for its interaction with ECM(55), and has been linked to other functions such as immune cell extravasation, response against pathogens, development of fibrosis, and wound healing(56, 57). This interaction is in line with the findings of our pathway analysis of genes differentially expressed in B cells from DCM and controls, that showed upregulation in leukocyte extravasation signaling, in both plasma cells and naive B cells (Figure 4). It is also in line with published evidence indicating that murine heart failure-derived B cells can cause increased fibroblast proliferation and collagen production(12), that CD44 activation on B cells can increase pro-inflammatory gene expression(58), and that B cells contribute to myocardial fibrosis in specific murine models of cardiomyopathy(59, 60).

Our analysis of cell communication in DCM highlighted two unexpected interactions that have not been described before: communication between B cells and eosinophils and communication between B cells and pericytes. We found predicted interactions from eosinophils to B cells that occurred through the MIF–CD44/CD74 complex (Figure 3B), which has been associated with mediating B cell chemotaxis and survival(38–40). The CD44/CD74 complex activates also NFκB signaling (61), which was an upregulated pathway in DCM plasma cells (Figure 4). However, eosinophils were also found to interact with B cells through galectin 9 – CD45 or CD44 (Figure 3B), an interaction that has been shown to inhibit B cell signaling and activation(41, 42). This suggests that eosinophils might contribute to the fine-tuning of B cell activation in DCM. In addition, we found that B cells signaled to eosinophils through laminin–CD44 and HLA- DRB1– CD4 (Figure 3A), which are strong activating receptors. This suggests that B cells may play a role in eosinophil activation in DCM(37, 62). Eosinophils have been shown to play a key pathogenic role in certain forms of DCM(63, 64), and thus this observation further corroborates the notion that B cells might play an important role in the pathogenesis of DCM. We found also a predicted interaction between B cells and pericytes (Figure 2 E-F and Figure 3). Pericyte THBS4 signaling to CD47 on B cells was uniquely seen in DCM patients. Notably, pericytes have been strongly associated with fibrosis in various disease states(65–67) and *Thbs4* has been identified as a key regulator of cardiac fibrosis in animal models(68). This suggests that myocardial B cells might play a role in DCM-associated cardiac fibrosis.

The analysis of genes differentially expressed in B cells between DCM and controls or ARVC and controls strengthens and expands the findings from our cell-cell interaction analysis. We found dysregulation of several pathways related to immune activation in DCM, in both naive and plasma cells (Figure 4 and Figure S3), that corroborate findings from murine models and from initial observations in humans. More specifically, the top dysregulated KEGG pathway in naive B cells from DCM patients was “antigen processing and presentation” (Table 3). “Antigen processing and presentation” was previously highlighted as one of the key pathways dysregulated in myocardial B cells in the context of murine post-ischemic dilated cardiomyopathy(10). DCM was characterized by metabolic activation of plasma cells (i.e. upregulation of oxidative phosphorylation pathway, Figure 4B and Table 4). Several studies have shown the presence of autoantibodies against cardiac proteins in DCM patients (69–71) and treatment with immunoglobulin adsorption has shown potential benefits for patients with myocardial autoantibodies (72–74). In the only clinical study that addressed the role of B cell depletion in DCM, antibody-mediated B cell depletion resulted in a marked clinical improvement in selected patients with chronic myocardial inflammation that did not respond to standard treatments(75). All things considered, therefore, our differential gene expression analysis supports the hypothesis that myocardial-associated B cells play a role in human DCM and suggests that the production of pathogenic antibodies in DCM might take place within the myocardium.

Our analysis is the first of its kind and provides several novel insights into the biology of human myocardial B cells, but it has several limitations that should be kept in mind when considering our findings. First of all, we integrated multiple, previously collected datasets. Therefore, we cannot exclude that “batch effects” might have biased our findings or that biologically important signaling pathways might have been missed due to the integration process. Second, while we integrated a large number of datasets corresponding to 45 normal human hearts and 70 hearts with DCM, we had data from only 8 patients with ARVC. Therefore, we cannot exclude that imbalances in sample size between DCM and ARVC might have reduced our statistical power in the analysis of ARVC patients. In addition, cell-cell interaction probabilities in scRNAseq and snRNAseq are based on expression levels and cell densities, but cannot take into account the spatial proximity of the cells. Additional work using spatial analyses will be necessary to better characterize the predicted interactions that we describe.

In summary, through the analysis of integrated single-cell datasets, we provide insights into the unique, dynamic, biological niche that B cells occupy within the human myocardium. Our findings provide novel insight into the biology of human myocardial B cells, corroborate previous work in murine models and clinical datasets, and support the notion that B cells play an important role in dilated cardiomyopathy. Further experimental work will be needed to confirm the extensive network of intracardiac intercellular communications that we describe and to further characterize B cell function in various forms of cardiomyopathy.

## ACKNOWLEDGMENTS

This publication was made possible by an NIH-funded postdoctoral fellowship to J.L. (T32-HL007227). The content of this manuscript has been published online as a pre-print in BioxRiv(76).

## AUTHOR CONTRIBUTIONS

Kevin C. Bermea and Carolina Duque led the data analysis, figure design, and manuscript writing. Aashik Bhalodia assisted with figure design and manuscript writing. Charles D. Cohen, Sylvie Rousseau, Jana Lovell, Marcelle Dina Zita, and Monica R. Mugnier contributed to the manuscript writing. Luigi Adamo coordinated the project and participated in writing and editing the manuscript. All authors approved the final version.

## DESCRIPTION OF SUPPLEMENTARY TABLES

**Supplementary Table 1.** List of biomarkers used for ScType cell classification.

**Supplementary Table 2.** DESeq statistical analysis of plasma cells derived from DCM patients versus controls.

**Supplementary Table 3.** DESeq statistical analysis of naive B cells derived from DCM patients versus controls.

**Supplementary Table 4.** DESeq statistical analysis of plasma cells derived from ARVC patients versus controls.

**Supplementary Table 5.** DESeq statistical analysis of plasma cells derived from ARVC patients versus controls.

**Supplementary Table 6.** Dysregulated GO pathways in ARVC’s plasma cells cluster, contrasting ARVC versus control samples data.

**Supplementary Table 7.** Dysregulated IPA pathways in ARVC’s naive B cells cluster, contrasting ARVC versus control samples data.

**Supplementary Table 8.** Dysregulated IPA pathways in ARVC’s plasma cells cluster, contrasting ARVC versus control samples data.

**Supplementary Table 9.** Dysregulated IPA pathways in DCM’s naive B cells cluster, contrasting ARVC versus control samples data.

**Supplementary Table 10.** Dysregulated GO pathways in DCM’s naive B cells cluster, contrasting ARVC versus control samples data.

**Supplementary Table 11.** Dysregulated IPA pathways in DCM’s plasma cells cluster, contrasting ARVC versus control samples data.

**Supplementary Table 12.** Dysregulated GO pathways in DCM’s plasma cells cluster, contrasting ARVC versus control samples data.

---

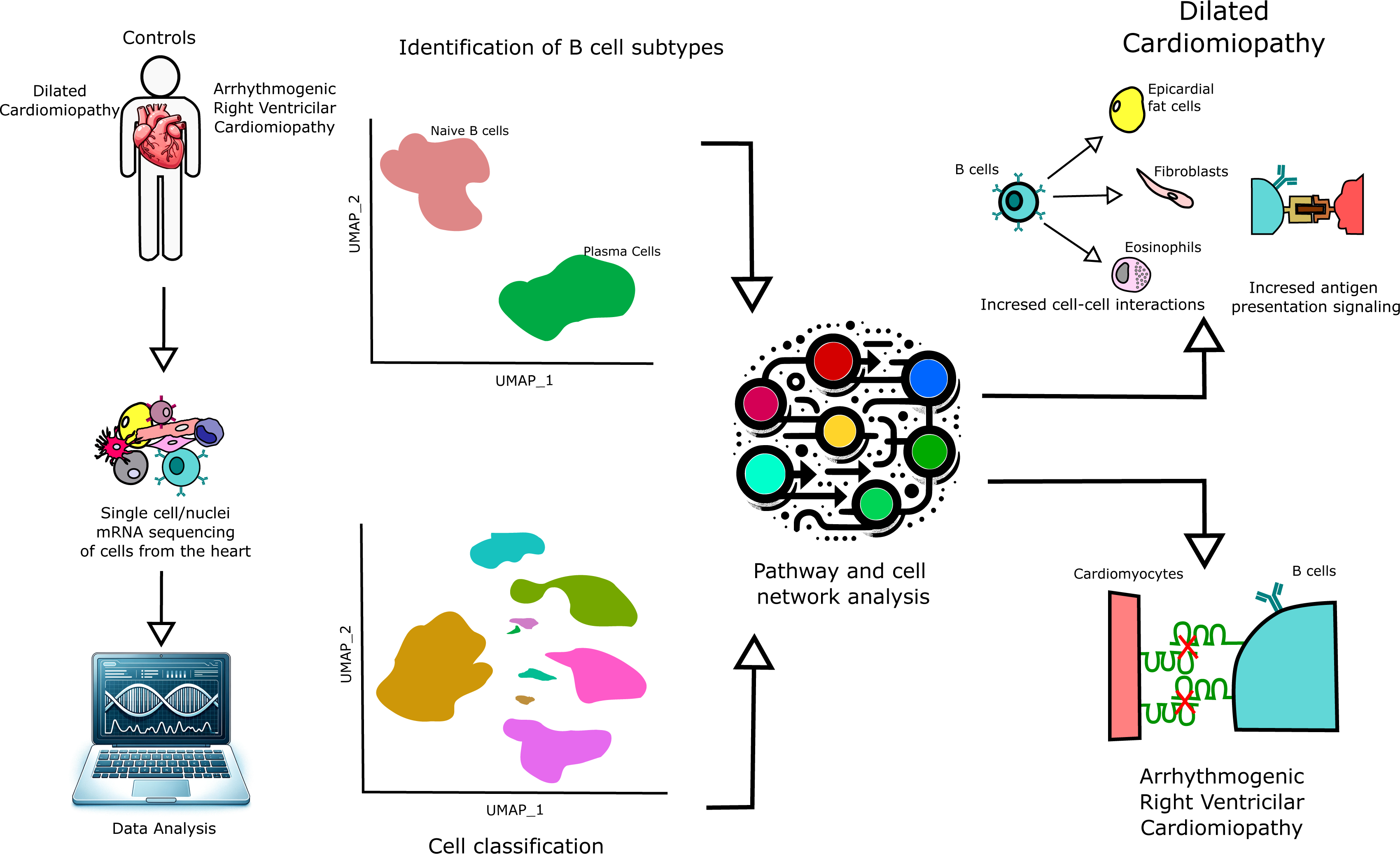

