## Supplementary material for "Myocardial B cells have specific gene expression and predicted interactions in Dilated Cardiomyopathy and Arrhythmogenic Right Ventricular Cardiomyopathy": Figure S3

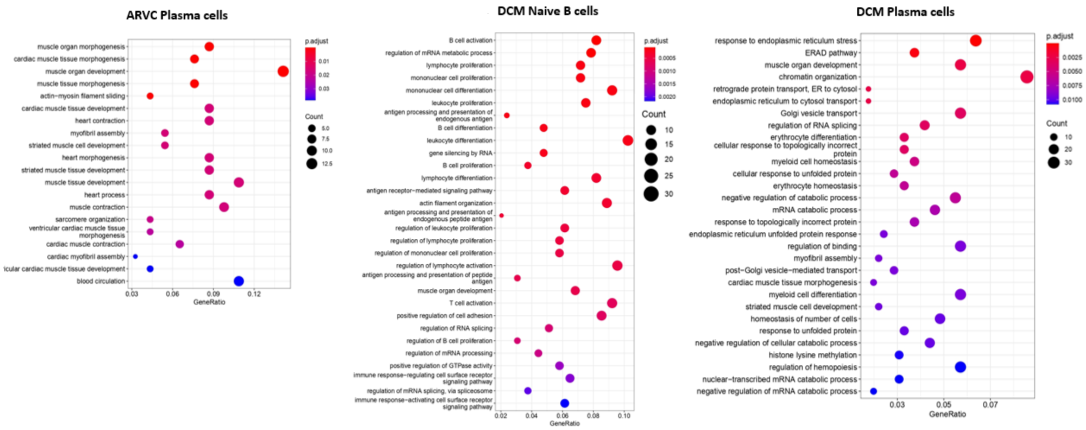


**Supplementary Figure 3. Gene ontology enrichment analysis of Naive and Plasma B cells in DCM or ARVC cardiac samples vs control**. a) Dysregulated gene ontology pathways in Plasma cells from ARVC samples. – No dysregulated gene ontology pathways were identified when comparing naive B cells from ARVC vs controls. b) Dysregulated gene ontology pathways in Naive B cells from DCM samples. c) Dysregulated gene ontology pathways in Plasma cells from DCM samples. DEGs with p-value < 0.05 calculated using DESeq2 and absolute fold change >1.5 were used for this analysis.
