## Supplementary material for "Myocardial B cells have specific gene expression and predicted interactions in Dilated Cardiomyopathy and Arrhythmogenic Right Ventricular Cardiomyopathy": Figure S2

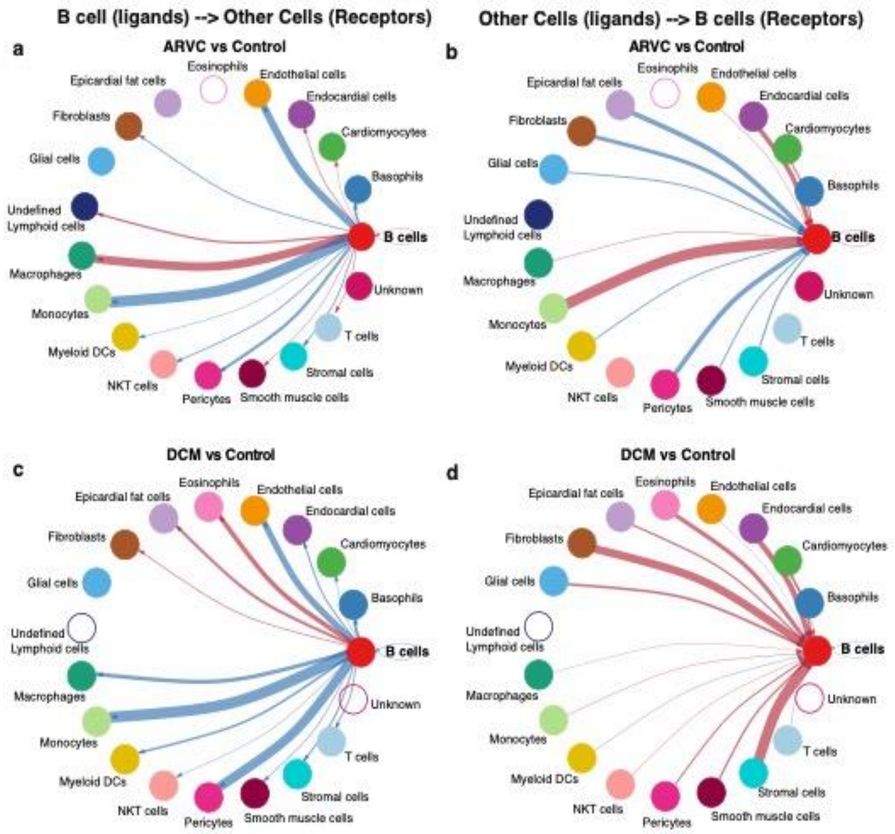


**Supplementary Figure 2. Circle plot of B cell interaction as compared to control.** The difference of interaction strength of B cell ligands to other cell's receptors between a) ARVC and controls and c) DCM and controls, and difference of interaction strength of other cell ligands to B cell receptors between b) ARVC and controls and d) DCM and controls are depicted. Thickness of the line is relative to the maximum difference in communication probability between disease conditions. Blue line color represents decreased interaction strength, red represents increased interaction strength. Open circles represent cell types that are not detected in either of the disease states being compared. DC = dendritic cell.
