## Supplementary material for "Myocardial B cells have specific gene expression and predicted interactions in Dilated Cardiomyopathy and Arrhythmogenic Right Ventricular Cardiomyopathy": Figure S1

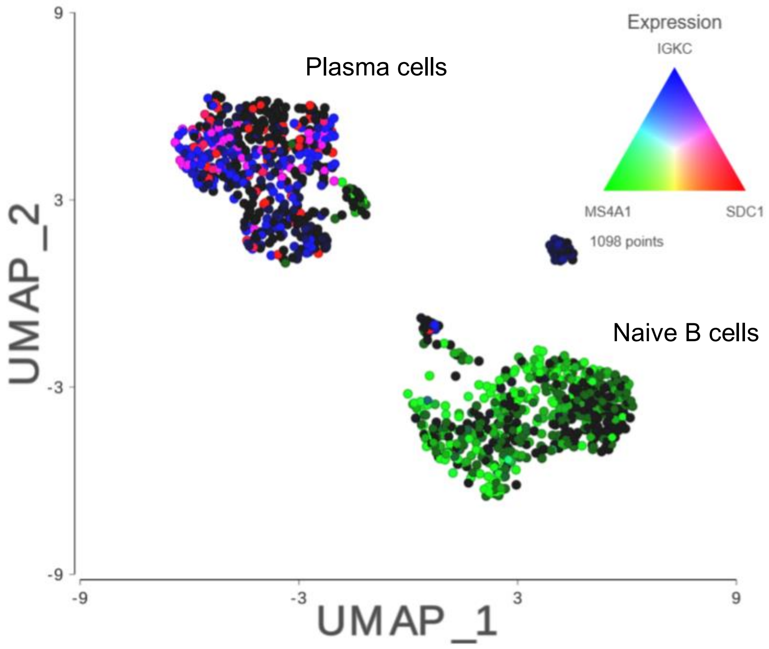


**Supplementary Figure 1. UMAP of the integrated and classified B cells from all datasets showing the markers for naive B cells and plasma cells.** B cells were subsetted from the integrated data and sub-classified into B- cell subtypes. The colors represent the expression levels of MS4A1 (CD20, green) highlighting the naive B cells; SDC1 (CD138, red); and IGKC (Ig Kappa Chain C, blue), highlighting the plasma cells. Data were normalized, transformed, and scaled using SCTransform with 3000 variable features, and ScType was used for cell classification.
